# Mapping multiple RNA modifications simultaneously by proximity barcode sequencing

**DOI:** 10.1101/2024.10.09.617509

**Authors:** Erdem Sendinc, Hua Yu, Yu Hsien Hwang Fu, Jerome Santos, Zachary Johnson, Justin R. Kirstein, Jiani Niu, Michael B. Chabot, Vito Adrian Cantu, Željko Džakula, Quan Lam, Ananya Anmangandla, Timothy S. Burcham, Eric M. Davis, Zachary D. Miles, Andrew D. Price, Byron W. Purse, Richard I. Gregory, Gudrun Stengel

**Author notes:** Corresponding authors: Richard I. Gregory, Gudrun Stengel.

## Abstract

RNA is subject to a multitude of different chemical modifications that collectively represent the epitranscriptome. Individual RNA modifications including N6-methyladenosine (m^6^A) on mRNA play essential roles in the posttranscriptional control of gene expression. Recent technological advances have enabled the transcriptome-wide mapping of certain RNA modifications, to reveal their broad relevance and characteristic distribution patterns. However, convenient methods that enable the simultaneous mapping of multiple different RNA marks within the same sample are generally lacking. Here we present EpiPlex RNA modification profiling, a bead-based proximity barcoding assay with sequencing readout that expands the scope of molecular recognition-based RNA modification detection to multiple targets, while providing relative quantification and enabling low RNA input. Measuring signal intensity against spike-in controls provides relative quantification, indicative of the RNA mod abundance at each locus. We report on changes in the modification status of HEK293T cells upon treatment with pharmacological inhibitors separately targeting METTL3, the dominant m^6^A writer enzyme, and the EIF4A3 component of the exon junction complex (EJC). The treatments resulted in decreased or increased m^6^A levels, respectively, without effect on inosine levels. Inhibiting the helicase activity of EIF4A3 and EIF4A3 knockdown both cause a significant increase of m^6^A sites near exon junctions, consistent with the previously reported role of EIF4A3 in shaping the m^6^A landscape. Thus, EpiPlex offers a reliable and convenient method for simultaneous mapping of multiple RNA modifications to facilitate epitranscriptome studies.

## Introduction

N6-methyladenosine (m^6^A) is the most widespread RNA modification on mRNA, present at roughly 0.2– 0.4% of all adenosines. The modification has been implicated in important diseases, such as various forms of cancer, neurological, cardiovascular and metabolic disorders. Today, the distribution of m^6^A along the mRNA transcripts (“m^6^A landscape”) is well established: m^6^A is situated in a consensus RNA DRACH motif (D = A/G/U, R = A/G, H = A/C/U) and is highly enriched in the 3’ untranslated region (UTR) (Dominissini et al., 2012; Meyer et al., 2012). METTL3, the core methyltransferase (MTase) of the m^6^A writer complex, specifically recognizes and binds to the DRACH motif (J. Liu et al., 2014; Wang et al., 2016). Other components of the m^6^A writer complex include METTL14, which forms a heterodimer with METTL3 and is essential for the MTase activity. Additional interactions with WTAP, VIRMA, RBM15/RBM15B, and ZC3H13 have been reported (Knuckles et al., 2018; Lence et al., 2016; Su et al., 2022). Even though DRACH motifs occur across the entire mRNA sequence, m^6^A sites are highly enriched in the 3’UTR, suggesting the existence of a mechanism beyond sequence-specificity that directs adenosine methylation to the 3’UTR (Dominissini et al., 2012; B. Linder et al., 2015). Recent studies reported a crucial role of EIF4A3 in shaping the m^6^A landscape (He et al., 2023; Uzonyi et al., 2023; Yang et al., 2022). EIF4A3 is an ATP dependent RNA helicase which forms the multimeric Exon Junction Complex (EJC) together with MAGOH, Y14, and MLN51. Depletion of EIF4A3 causes increased m^6^A methylation of DRACH motifs of short internal exons, and sites close to exon-exon junctions within mRNA (He et al., 2023; Uzonyi et al., 2023; Yang et al., 2022). This suggests a model where the m^6^A landscape is shaped by the EJC protecting nascent transcripts from the m^6^A MTase machinery (He & He, 2024). This process results in absence of m^6^A from short exons and leads to concentration of m^6^A at the 3’UTRs that are typically devoid of introns and therefore unprotected by the EJC. Abnormal expression or mutations of EIF4A3 have been associated with various types of cancer, suggesting that the increased m^6^A levels observed in many cancers may be a result of aberrant splicing machinery and incomplete protection of exon junctions by the EJC.

Another abundant modification on nascent RNA pol II transcripts is A-to-I (adenosine-to-inosine) RNA editing by ADAR (adenosine deaminase acting on RNA) enzymes (Savva et al., 2012; Tan et al., 2017). Inosine typically occurs in double-stranded RNA (dsRNA) regions, which can form through intramolecular base-pairing within a single RNA molecule or between two separate RNA molecules. ADAR enzymes recognize these dsRNA structures, bind them, and catalyze the deamination of adenosine’s amine group to generate inosine. A-to-I editing is known to increase the diversity of coding sequences, modulate the immune response, influence RNA splicing, stability, and interactions with RNA-binding proteins, thus regulating gene expression (Jarmoskaite & Billy Li, 2024; Keegan et al., 2023; Savva et al., 2012; Shiromoto et al., 2021; Song et al., 2022; Tan et al., 2017; Tang et al., 2020). Dysregulation of A-to-I RNA editing has been associated with various diseases, including neurological disorders, autoimmune diseases, and cancers (Song et al., 2022). Multiple companies are currently working on therapeutic RNA editing approaches that exploit antisense oligos in conjunction with endogenous ADAR activity to correct mutations by means of codon editing (Booth et al., 2023; Mullard, 2024). Even though treating genetic diseases by RNA editing requires lifelong treatment, it has fewer side effects and does not bear the risk of irreversible, permanent genome alterations.

With growing recognition of the vital role of RNA modifications in gene expression, many sequence-resolved protocols for the detection of RNA modifications have been published. The most common RNA modifications are N6-methyladenosine (m^6^A), inosine (I), and pseudouridine (Ψ), which are present in both coding and non-coding RNAs. Rarer RNA modifications, such as N5-methylcytosine (m^5^C), N1-methyladenosine (m^1^A), and N7-methylguanosine (m^7^G) have also been characterized and mapped across transcriptomes(Zaccara et al., 2019). RNA modification detection and genome-wide mapping methods include immunoprecipitation of methylated RNA by antibodies (meRIP) and a broad range of conversion methods where treating RNA with a modification-specific chemical or enzyme generates a signal in cDNA, such as strand truncation, base deletion, or a mutational hotspot. GLORI and eTAM-Seq are methods that chemically or enzymatically convert all unmethylated A to inosine (I), which is read as G, allowing m^6^A detection at single nucleotide resolution (C. Liu et al., 2022; Xiao et al., 2023). While chemical conversion methods have the advantage of providing single base resolved localization of the modification, they tend to be destructive to RNA and require high RNA inputs.

The methods above are designed for detecting a specific RNA modification and are inherently incapable of multiplexed detection. Owing to the lack of multiplexed, sequence resolved epitranscriptomic methods, very little is known about correlative, quantitative changes in multiple RNA modifications in response to—or as a driver of—different molecular mechanisms. Even without sequence resolution, mass spectrometry studies have reported changes in the total epitranscriptomic landscape in cancer with predictive value for tumor classification(Sun et al., 2019). While direct sequencing of RNA with nanopores holds promise for detecting multiple RNA modifications concurrently with single base resolution and strand information, the method is still limited by high error rates and the need for an unmodified reference, training set and machine learning for data interpretation (H. Liu et al., 2019).

Despite the large number of published methods, most studies of m^6^A in biological systems have used m^6^A immunoprecipitation (meRIP) and variants thereof (Dominissini et al., 2012; Meyer et al., 2012). While all methods have strengths and weaknesses that must be considered for a particular use case, we find that there are three prominent barriers to adoption of newer methods: (i) high RNA input requirements that limit studies to RNA extracted from immortalized cell lines, (ii) lack of commercial reagents for detection of multiple RNA modifications simultaneously, and (iii) the availability of bioinformatics tools to analyze datasets, which is particularly challenging in nanopore sequencing.

Here, we present the EpiPlex proximity barcoding assay for the multiplexed detection of m^6^A and inosine, with demonstrated scalability to additional RNA modifications. The assay requires 20 ng of coding or 250 ng of total RNA as an input, provides quantitative metrics on RNA mod abundance and is commercially available as a reagent kit including a data analysis pipeline. In this study, we demonstrate the utility of the EpiPlex assay in an investigation of the m^6^A methylation and A-to-I editing of mRNAs regulated by METTL3, EIF4A3, and ADAR enzymes.

## Results

### EpiPlex Assay Architecture and Workflow

The foundation of the EpiPlex assay is a pool of magnetic beads that display multiple copies of RNA-modification specific protein binders and barcoded Illumina adapters. We term the beads “monoclonal beads” because each bead type exhibits only a single type of binder and adapter at an optimized ratio. In addition to universal sites for PCR amplification, the adapter sequence comprises a modification-specific barcode (MBC) and a unique molecular identifier (UMI) **(FIG. 1)**. Including a UMI allows for the correction of PCR errors, enabling accurate deduplication of sequencing reads. The surface density of molecules is such that the adapter can be incorporated into the cDNA in a process we call proximity barcoding by reverse transcription. Transferring the adapter in this way records the identity of the RNA modification that was captured by the bead. The surface architecture and barcoding reaction have been carefully optimized to occur intra-bead, which is essential for preventing off-target barcoding.

**FIG. 1:**
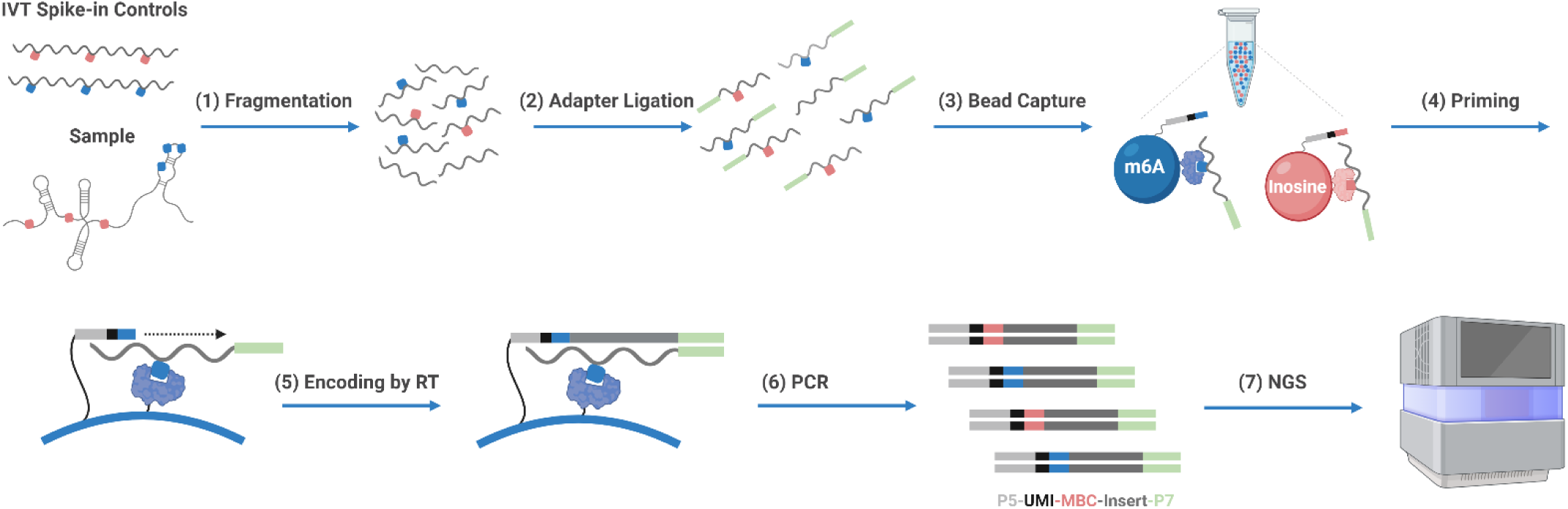
Schematic of the multiplexed EpiPlex barcoding workflow. RNA is mixed with spike-in controls, fragmented, and captured by a pool of two bead types, where each bead type exhibits a distinct RNA modification barcode and protein binder. The barcode is incorporated to the cDNA by reverse transcription and the library is completed by PCR amplification. Sequencing infers the presence of an RNA modification based on the barcode. The resulting library architecture comprises the Illumina P5 adapter (light grey), a Unique Molecular Identifier (UMI; black), a Modification Barcode (MBC; blue for m^6^A, red for inosine) and the Illumina P7 adapter (light green). RNA is displayed as wiggly lines, DNA as straight lines.

The total library preparation workflow encompasses the following steps (**FIG. 1**): Purified RNA is mixed with spike-in controls, fragmented to ∼ 140bp, end-repaired and ligated to the P7 adapter (1). RNA fragments are captured by specific RNA modification binders (2). The captured RNA fragments are random primed by bead-anchored P5 adapters (3). The immobilized P5 adapters comprise the MBC, which is transferred to the cDNA during reverse transcription. PCR amplification concludes the library preparation (5) and paired-end sequencing is used to read the UMI, MBC along with the RNA sequence. In total, the workflow takes 7 hours with 3 hours of hands-on time. To account for the dynamic range of the RNA expression levels, the reaction is split after step (1) into an “enrichment reaction” mixture, which is exposed to the beads and barcoded in an RNA-modification specific manner, and a “solution control”, which randomly barcodes all RNA fragments – modified or not.

While researchers often use abundant RNA isolated from cells in culture for mechanistic studies, EpiPlex methodology is also applicable to samples with low amount of input RNA. Tissue samples, clinical biopsies, and other samples of native biological material typically provide only small amounts of RNA, often containing less than 500 ng of total RNA. mRNA makes up typically 1–3% of this total transcriptome. In this study, we used 50 ng of polyA-selected RNA input and we validated the assay down to 20 ng of poly-A RNA and 250 ng of total RNA.

### The impact of METTL3 inhibition and METTL3 knockdown

To test the EpiPlex platform, we first mapped m^6^A modifications globally in HEK293T cells under different conditions. This new methodology yielded m^6^A peaks which highly overlapped with the peaks obtained by existing m^6^A mapping techniques, such as GLORI and meRIP-seq, validating our new methodology (**SI FIG. 1**). We subsequently assessed the technical reproducibility of EpiPlex by analyzing the same sample on different days. (**SI FIG. 2A**). This led to highly consistent numbers of m^6^A peaks, with percent standard deviations at 0.80%.

To test the dynamic range of EpiPlex, we decreased the global m^6^A levels using two approaches: in one approach, we treated HEK293T cells with the METTL3 inhibitor STM2457 for two days and subjected the polyA-selected RNA to the EpiPlex assay (Yankova et al., 2021). In the other approach, we utilized a HEK293T cell line where the endogenous METTL3 gene was replaced by an N-terminally degron-tagged METTL3 (Nabet et al., 2018). Treatment of these cells with the degrader molecule (dTAG) rapidly leads to degradation of METTL3 enzyme, which results in a loss of *de novo* m^6^A detected by RNA mass spectrometry (**SI Fig 3A and 3B**). We analyzed RNA isolated from either METTL3 inhibitor or degrader treated cells employing EpiPlex. In both cases, the EpiPlex methodology identified a significant decrease in m^6^A, as evidenced by a sharp decrease in the number of m^6^A peaks (**FIG. 2A**). While most transcripts lost m^6^A upon degradation or enzymatic inhibition of METTL3, some m^6^A loci remained methylated (**FIG. 2A**). Upon further investigation, EpiPlex revealed that these persistent loci display METTL3 methylation motif (DRACH) with a diminished m^6^A enrichment, suggesting that these loci might be a result of remaining METTL3 activity due to incomplete METTL3 degradation or inhibition and/or present on mRNAs with a long half-life (**FIG 2B**). A representative gene track of m^6^A enrichment is shown in **FIG. 2B**. Enrichment values were obtained by normalizing the raw data to the spike-in and to the solution controls as explained in the supporting info (**SI Figs 4-6**). Taken together, EpiPlex mapping methodology detected m^6^A quantitatively and revealed its global distribution. Additionally, it enabled detection of changes in m^6^A level upon disruption of the MTase machinery by two independent methods.

**FIG. 2:**
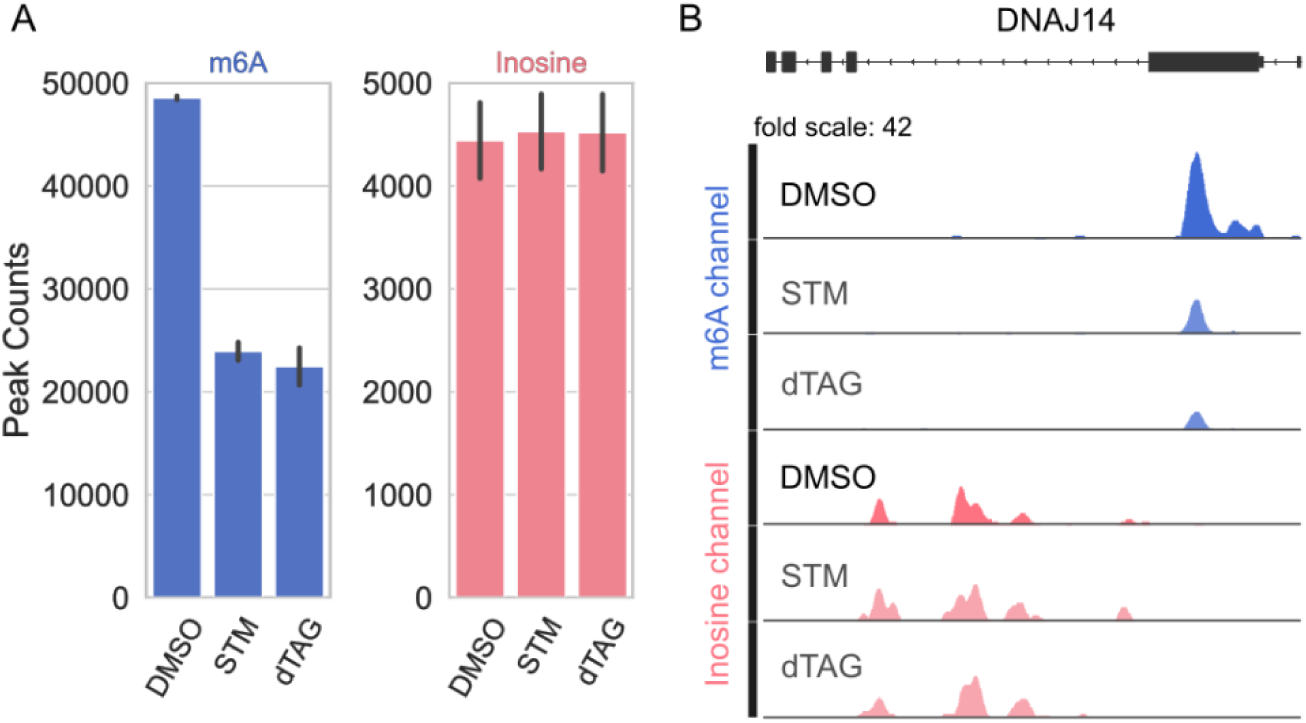
m^6^A methylation decreases upon METTL3 inhibition or degradation. **(A)** EpiPlex analysis of global number of m^6^A (blue) and inosine (red) peaks upon STM2457 inhibitor (STM) or dTAG degrader treatment, compared to the DMSO control (DMSO). The error bars indicate the standard deviation of three technical replicates per condition. **(B)** m^6^A and inosine enrichment data of the DNAJ14 gene transcript upon STM2457 inhibitor or dTAG degrader treatment. The gene tracks illustrate a decrease in m^6^A enrichment in the treated conditions, while the inosine levels remain largely unchanged.

EpiPlex enables simultaneous investigation of multiple RNA modifications. Beyond mapping m^6^A, we used this methodology to identify inosines resulting from A-to-I editing of the RNA transcriptome. By allowing us to map both m^6^A and inosines from the same RNA sample, EpiPlex facilitated our investigation into how the disruption of m^6^A machinery affects another co-transcriptional modification, A-to-I editing. EpiPlex allows detection of A-to-I editing within the RNA samples by first enriching inosine containing RNAs with inosine capturing monoclonal beads (**FIG. 1**). A-to-I editing is detected when inosine is read as guanine during sequencing of bead captured RNA molecules containing modification specific adapters, leading to an of A-to-G mutation under each inosine enrichment peak. This enables mapping of this RNA modification globally with single base resolution. Despite the sharp decrease in m^6^A, neither the loss of METTL3 protein nor the loss of METTL3 activity resulted in a significant change in global inosine peak number, suggesting that ADAR mediated inosine machinery is mechanistically independent from METTL3 mediated m^6^A methylation (**FIG. 2A and 2B**). This is in line with the observation that METTL3 acts on DRACH motifs in single-stranded (ss) RNA (Dominissini et al., 2012; Meyer et al., 2012), whereas ADAR activity targets regions of double-stranded (ds) RNA without distinct sequence preference, and these enzymes are not known to interact with one another.

### Quantitative Detection of METTL3 Inhibition

To determine whether EpiPlex can detect the changes in RNA modifications quantitatively, we titrated cultured HEK293T cells with increasing concentrations of the METTL3 inhibitor STM2457 (Yankova et al., 2021). Before peak calling, the data set was downsampled to equal read coverage at the MBC level and transformed, including normalization to the spike-in and solution controls and smoothing was applied (**SI FIGs. 4 and 5**). The polyA-enriched RNA of unperturbed control cells exhibits ∼53,000 m^6^A peaks and ∼3,500 inosine peaks. Upon inhibitor treatment, the number of m^6^A peaks decreases to ∼10,000, whereas the number of inosine peaks remains unchanged with minor noise fluctuations (**FIG. 3A**). Note that the inosine signal is inherently noisier due to the lower read coverage. In this experiment, m^6^A MBC reads were covered at 35M reads and inosine MBC reads at 1.5M. In accordance with the literature, the m^6^A methylation is enriched around the 3′-UTR of polyA RNAs (**FIG. 3B, blue lines**) and a significant enrichment of DRACH motif is observed under m^6^A peaks (**FIG. 3C, blue lines**), validating the accuracy of m^6^A identification (Sendinc & Shi, 2023). Although there is a significant drop in m^6^A methylation upon inhibitor treatment (**FIG. 3A and 3D**), the remaining m^6^A distribution still displays 3′-UTR enrichment and retains the DRACH motif (**FIG. 3B and 3C**), suggesting that the inhibitor does not impact the m^6^A location, although METTL3 is inhibited.

**FIG. 3:**
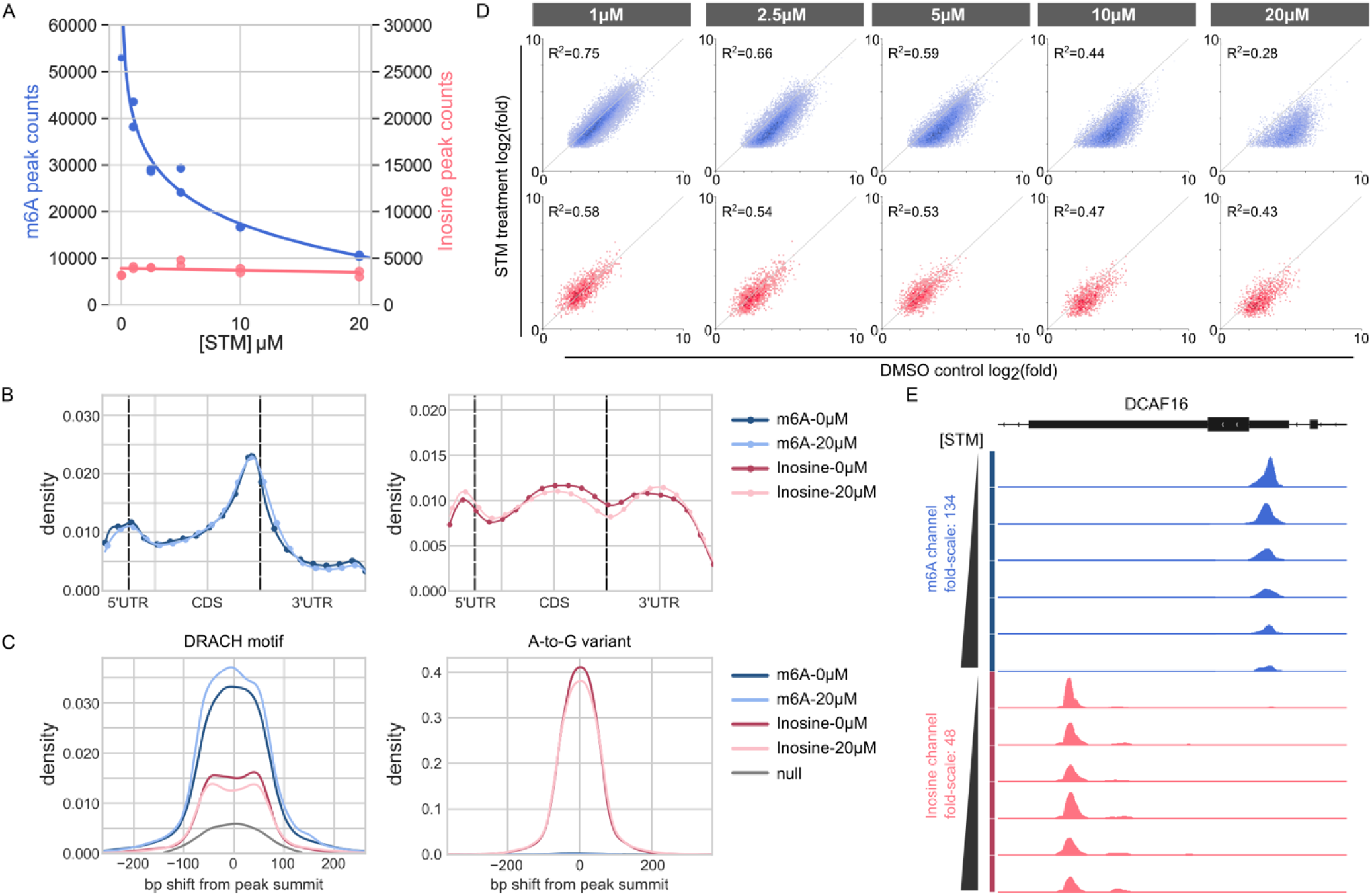
Titration of HEK293T cells with METTL3 Inhibitor. **(A)** Global number of m^6^A (blue) and inosine (pink) peaks at different STM2457 inhibitor (STM) concentrations. **(B)** Transcript distributions of RNA modifications (m^6^A in shades of blue, inosine in shades of red). m^6^A peaks are enriched at the 3’ untranslated region (3′-UTR) consistent with previous reports, inosine exists in both UTRs and in the intron regions of the coding sequence (CDS). For both RNA mods, the distribution profile is the same for untreated (“0uM”) and inhibitor treated cells (“20uM”). **(C)** Motif shift plots. Enrichment and distance of the DRACH motif from peak summits and enrichment and distance of A-to-G mutations from peak summits. Motif shifts under m^6^A peaks are shown in shades of blue, motif shifts under inosine peaks in shades of red. The gray curve describes the null distribution of DRACH motifs in the transcriptome. Note that the total m6A peak numbers described by the transcript distribution plot (B) and motif density plot (C) are different +/- METTL3 inhibitor. The fact that neither plot changes +/- inhibitor treatment supports that the m6A peaks that remain after inhibitor treatment have the same characteristics – they are in the 3’UTR at the same location (despite lower modified transcript abundance) and have a similar width. Only the fold-enrichment (e.g. equivalent to the peak height in figure E) changes. **(D)** Differential analysis of the fold-enrichment values of m^6^A and inosine peaks of untreated versus treated cells. At low STM concentration, the data points are symmetrically distributed around the diagonal with R^2^ values of 0.75 and 0.58 for m^6^A and inosine, respectively. Adding higher concentrations of inhibitor shifts the m^6^A data points below the diagonal, indicative of a decrease in m^6^A abundance at all loci. **(E)** Gene viewer presentation of m^6^A and inosine sites in the DCAF16 gene. The EpiPlex assay produces gene tracks for each MBC. Shown are bigwig traces of the transformed data used for peak calling (m^6^A sequencing read pileups in blue, inosine pileups in red). The y-axis represents the fold-enrichment of the RNA mod at each locus for increasing concentration of inhibitor.

Compared to m^6^A distribution, inosine shows a different modification profile along the transcripts (**FIG. 3B, red lines**). It is enriched around the 3’ and 5’UTRs and in the unspliced introns of the coding sequence (CDS) region (in the transcript plot introns are counted as part of the CDS region). Because the input RNA was polyA-enriched, the intron reads likely stem from pre-mRNA and unspliced introns, which explains the lower read coverage. Compared to m^6^A, inosine is rare in the exons of the CDS, likely because A-to-I editing can alter codon reading and cause amino acid substitutions during ribosomal decoding (Licht et al., 2019). Despite the significant changes in m^6^A, the treatment of cells with various doses of METTL3 inhibitor did not alter the abundance or distribution of A-to-I editing, further indicating that this modification is mechanistically distinct from m^6^A methylation, although these modifications can co-occur on the same nascent mRNA molecule. Inhibitor treatment resulted in a dramatic drop in the number of m^6^A loci, but not a full elimination (**FIG. 2A and 3A**). Differential analysis of the fold-enrichment values, a quantitative metric of RNA modification abundance, revealed that even the remaining peaks all decreased in magnitude and share the METTL3 DRACH motif, consistent with a lower abundance at these m^6^A sites (e.g. **FIG. 2B**). Plotting the fold-enrichment values for all m^6^A sites of the control versus the same sites in the inhibitor-treated samples showed that the m^6^A enrichment decreased at all but ∼ 200 loci with increasing dose of the METTL3 inhibitor (**FIG. 3D).** Representative R^2^ values for the fold-enrichment of m^6^A and inosine peaks in technical replicates are shown in **SI FIG. 2B**. The gradual, concentration dependent decrease of m^6^A abundance upon inhibitor treatment is demonstrated for the DCAF16 gene, a gene that also features a prominent inosine site that is unaffected by the METTL3 inhibitor (**FIG. 3E).**

The relative quantification provided by the EpiPlex assay enables the presented dose-response curve analysis. In addition to improving the accuracy of comparisons within each experimental group, the relative quantification feature of the EpiPlex assay reduces variability between experiments. For example, we included the untreated DMSO controls in three independent experiments and observed excellent reproducibility with a CV (coefficient of variation calculated by dividing the standard deviation of the number of m^6^A peaks across experiments by its mean) of less than 2.5% (**SI FIG. 2A**).

### Regulation of RNA modification landscape by EIF4A3

We and others have recently determined that EJC plays an important mechanistic role in m^6^A deposition by METTL3 complex (He et al., 2023; Uzonyi et al., 2023; Yang et al., 2022). Knockdown of the RNA-binding subunit of the exon-junction complex EIF4A3 results in an increase in global m^6^A methylation. This increase specifically occurs within ∼200 nt of 5’ and 3’ of splice sites (Uzonyi et al., 2023; Yang et al., 2022). This supports a molecular model where EJC binding to sites at exon-exon boundaries locally protects transcripts from methylation by METTL3 (He & He, 2023). The core EIF4A3 subunit assembles into a mega-dalton-sized RNP that occupies 150-200 nt footprints and suggest that the RNA within these complexes is packaged into an overall compact mRNP structure (Singh et al., 2012). These higher order EJC mRNP particles could mechanistically protect mRNAs from nucleases and RNA modifications.

In addition to being an RNA binding protein, EIF4A3 is also an ATP-dependent DEAD-box helicase. Besides forming high affinity RNA clamps, DEAD box helicases traverse RNA during unwinding thereby interacting with longer stretches of RNA (P. Linder & Jankowsky, 2011). Indeed, EIF4A3 displays RNA helicase activity, which is enhanced by EJC member CASC3 *in vitro* (Noble & Song, 2007). We tested whether this enzymatic activity of EIF4A3 is required in regulation of m^6^A deposition. We used a small molecule inhibitor that inhibits both the ATPase and helicase activities of EIF4A3 but does not inhibit other closely related DEAD-box helicases *in vitro* (Iwatani-Yoshihara et al., 2017). This inhibitor was shown to impact other EIF4A3 dependent processes in cells, such as nonsense-mediated RNA decay (NMD) (Iwatani-Yoshihara et al., 2017; Li et al., 2022). Upon treatment of cells with this EIF4A3 inhibitor, we confirmed that the inhibitor did not impact total EIF4A3 protein level in treated cells (**SI Fig. 3C)**. We carried out EpiPlex mapping and observed the canonical m^6^A enrichment near 3’ UTR of RNA pol II transcripts (**FIG. 4B)**. Although this 3’ UTR m^6^A enrichment did not change upon EIF4A3 inhibition significantly, we observed *de novo* m^6^A methylation over exons of mRNAs upon loss of EIF4A3 activity (**FIG. 4B-F)**. We also mapped m^6^A after knocking down EIF4A3 via siRNA. This led to a similar increase in m^6^A methylation (**FIG. 4A)**. Both the loss of EIF4A3 protein via siRNA knock-down and inhibition of EIF4A3 activity with an inhibitor resulted in a similar increase in m^6^A specifically over exons of transcripts, without impacting the canonical 3’ UTR m^6^A significantly. This indicates that not just the EIF4A3 protein but also its enzymatic activity is required for regulation of m^6^A deposition on RNA pol II transcripts.

**FIG. 4:**
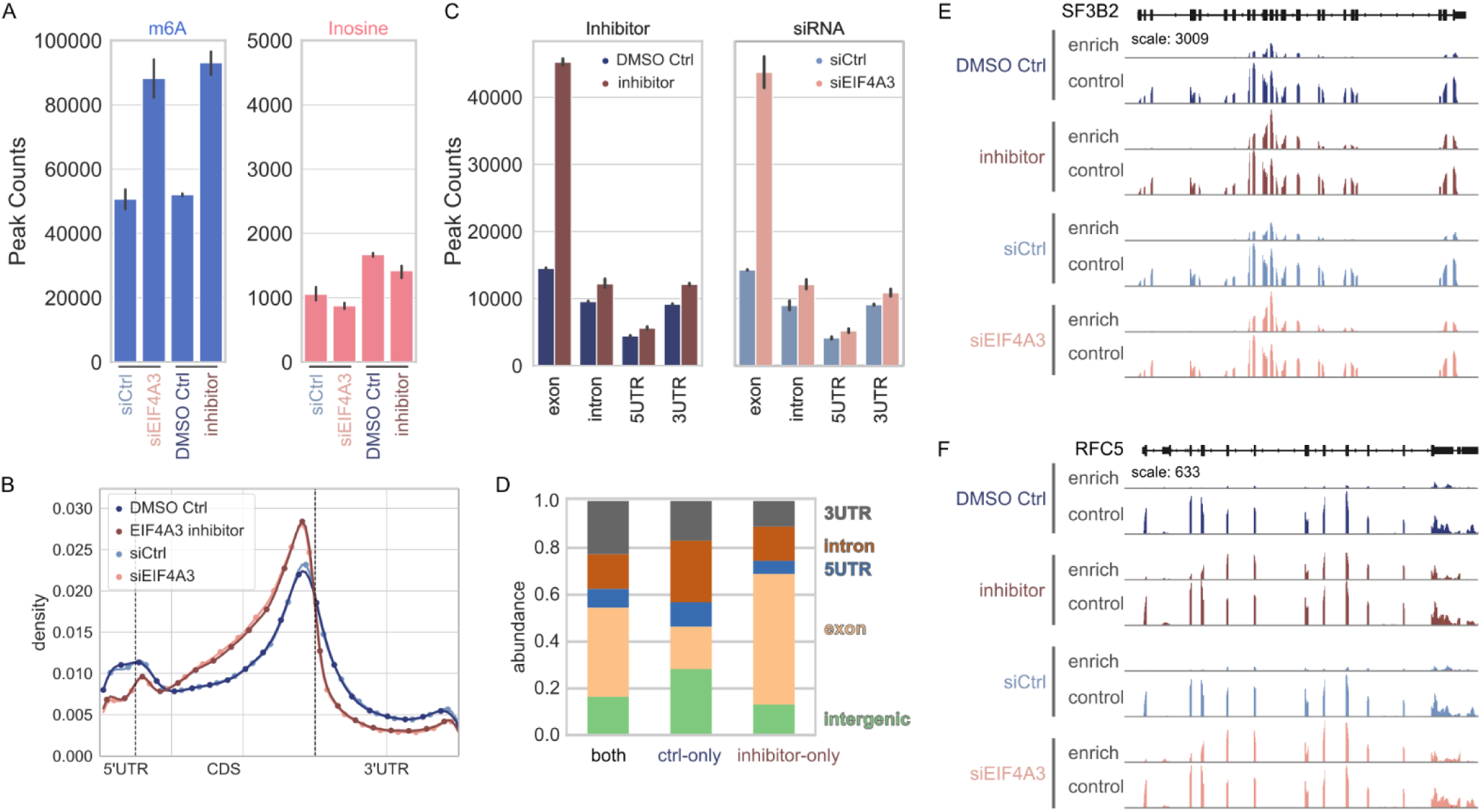
Effects of EIF4A3 inhibition on the landscape of the m^6^A epitranscriptome. **(A)** Inhibition of EIF4A3 by a small molecule inhibitor (DMSO Ctrl: untreated; inhibitor: treated) or by genetic knockdown (siCtrl: untreated; siEIF4A3: treated) leads to a global doubling of the number of m^6^A sites. Inosine levels are the same for untreated and treated samples. **(B)** Overlaid transcript distributions of m^6^A. Both untreated controls overlap (dark and light blue lines) and both treated samples overlap regardless of method of EIF4A3 inhibition (dark and light brown lines). The distribution of m^6^A peaks is notably shifted towards the CDS region for the inhibitor treated samples. **(C)** Absolute number of m^6^A peaks by gene feature. Regardless of the method of EIF4A3 inhibition, the new m^6^A marks are deposited in exon regions. **(D)** Relative abundance of m^6^A peaks in different gene regions. Comparison of m^6^A peaks that were found in untreated and treated samples, only in the untreated samples or only in the inhibitor treated samples. m^6^A peaks that are unique to the inhibitor sample reside mostly in the exons. **(E)** Gene viewer presentation of raw read pileups for the SF3B2 gene. The “enrich” gene track shows the reads obtained from the library that was exposed to the EpiPlex beads, whereas the “control” reads stem from the solution library. Neither untreated samples (DMSO Ctrl and siCtrl) show m^6^A signal in the exons of the “enrich” reaction; however, after treatment with inhibitor or EIF4A3 knockdown (inhibitor and siEIF4A3) m^6^A peaks appear in the same exon regions. **(F)** Gene viewer presentation of raw read pileups for the RFC5 gene [as in (E)].

Another abundant co-transcriptional RNA modification is A-to-I editing of RNA pol II transcripts carried out by ADAR enzymes. ADAR enzymes bind double-stranded RNA and enzymatically convert adenosines to inosines (Savva et al., 2012). Given that EIF4A3 is a transcriptional helicase, we tested whether its helicase unwinding activity might impact ADAR mediated dsRNA editing of RNA pol II transcripts. In addition to m^6^A methylation, EpiPlex mapping strategy allows us to determine A-to-I editing loci on mRNAs. We determined that neither the siRNA mediated knock down of EIF4A3 nor the inhibition of EIF4A3 activity impacted ADAR mediated A-to-I editing of mRNAs, significantly (**FIG. 4A)**. This suggests that effect of EIF4A3 inhibition is specific to METTL3 mediated m^6^A modification and does not impact other co-transcriptional process of A-to-I editing mediated by ADAR enzyme.

### Validation of Inosine signals by ADAR knockout

To investigate A-to-I editing of mRNAs with EpiPlex methodology further, we utilized ADAR1 knock-out HEK293T cells (**SI Fig. 3D**). Human genome encodes multiple adenosine deaminases acting on RNA (ADAR enzymes) (Savva et al., 2012). While ADAR1 (ADAR) and ADAR2 proteins can modify adenosines on RNA pol II transcripts, brain specific ADAR3 protein is catalytically inactive and has a regulatory role (Raghava Kurup et al., 2022). Human genome also encodes two other deaminases ADAT2 and catalytically inactive ADAT3, which are specific to tRNAs (Dolce et al., 2022). To investigate A-to-I editing of RNA pol II transcripts, we knocked out ADAR1 (both p150 and p110 isoforms) in HEK293T cells. We analyzed mRNA from these cells employing EpiPlex, which relies on A-to-G mutations for detection of inosines. Depending on the thresholding parameters set in our analysis pipeline and sequencing read depth, 80 to 98% of inosine peaks exhibit an A-to-G mutation. In the absence of ADAR enzyme, we observed an almost complete loss of inosines from mRNAs (a ∼10-fold drop in the global inosine levels) validating the accuracy of the EpiPlex assay (**FIG. 5A**). This also suggests that majority, if not all, mRNA A-to-I editing are dependent on ADAR1 enzyme in these cells.

**FIG. 5:**
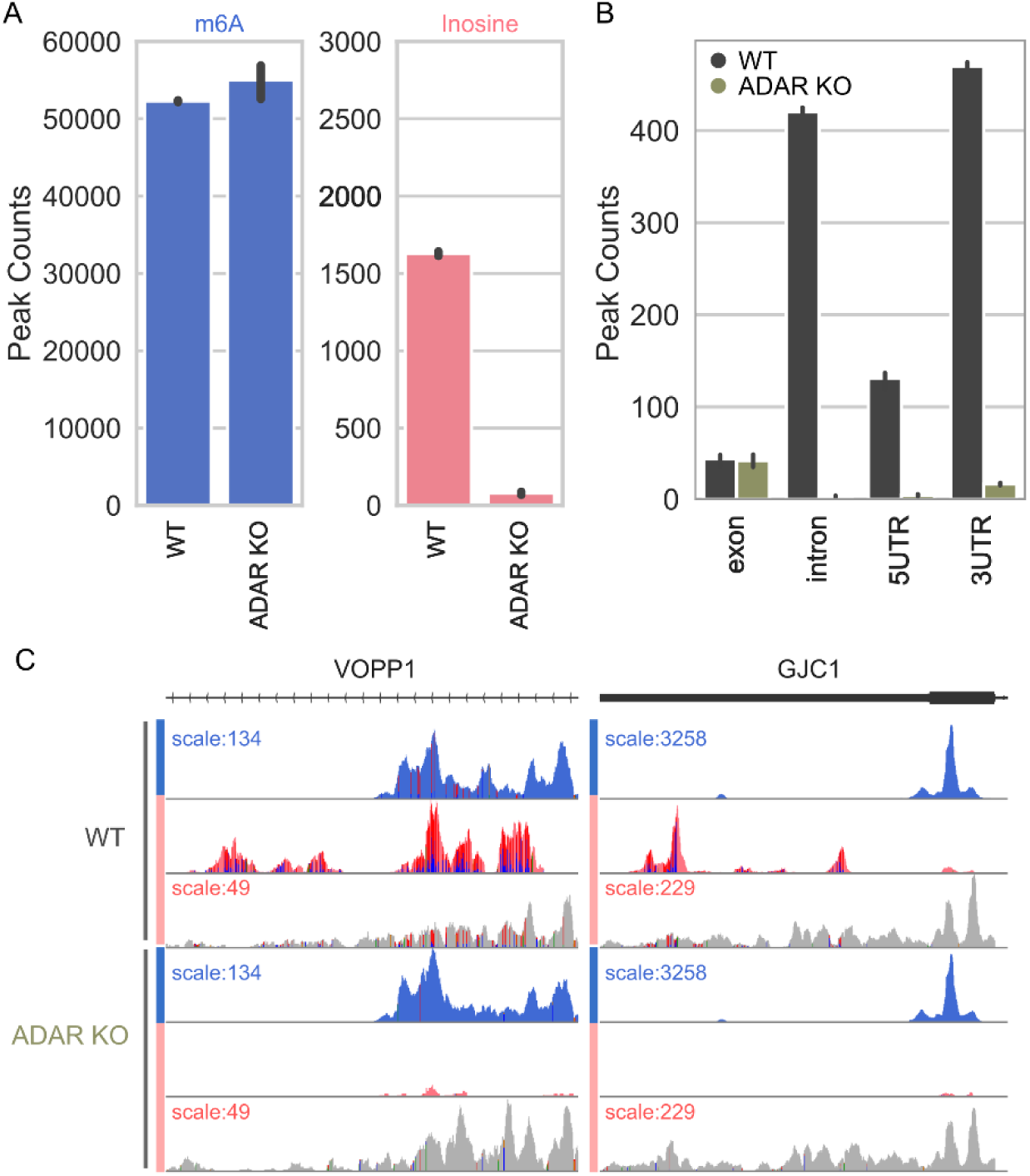
ADAR knockout. **(A)** Global number of m^6^A and inosine peaks in wildtype (WT) and ADAR knockout (ADAR KO) samples. The ADAR knockout eliminates most inosine peaks without affecting the m^6^A peaks. **(B)** In control cells, A-to-I edits are enriched in the introns, 5′-UTR and intergenic regions. Knockout of ADAR removes those signals nearly quantitatively, with minimal signal remaining in the exons and 3′-UTR. (**C**) Examples of raw read pile ups of a gene with inosine sites (pink) co-localized with m^6^A (blue) (c.f. VOPP1), and of a gene with inosine sites in the vicinity to m^6^A sites (c.f. GJC1). The vertical lines inside the inosine peaks indicate the A-to-G mutations. In the absence of ADAR, the inosine peaks disappear, while the m^6^A peaks persist, demonstrating the decoupling of RNA editing from m^6^A deposition and the independence of signal acquisition in the multiplex assay.

EpiPlex allows simultaneous analysis of m^6^A and A-to-I editing of the same RNA samples. This analysis revealed that the distribution of inosine and m^6^A are fundamentally different along the transcripts. While m^6^A accumulates in the 3′-UTR, inosine is found in all non-coding regions including the 3’UTR, 5’UTR and introns (**FIG. 3A and 5B**). Moreover, almost complete loss of A-to-I editing in ADAR knock out cells does not impact either m^6^A enrichment or the m^6^A distribution (**Fig 5A**). An example gene is depicted in Fig 5C, where the raw EpiPlex output with sequencing reads grouped according to their MBC in a genome browser. Raw read pileups are displayed to visualize the A-to-G mutations (vertical lines under peaks). Remarkably, the VOPP1 gene is an example of co-localization of m^6^A and inosine. The deposition of m^6^A methylation and inosine editing appear to be independent processes.

## Discussion

This study is the first demonstration of a proximity barcoding assay for the multiplexed detection of RNA modifications. Using a 2-plex detection scheme for m^6^A and inosine, we explored the possible interrelationship between these two mRNA modifications and deepened the understanding of the role of EIF4A3 in shaping the m^6^A epitranscriptome.

As previously reported, EIF4A3 is an essential part of the EJC and its presence is required for protecting short exons and exon junctions from adenosine methylation by METTL3 complex. This protective action results in m^6^A enrichment in the 3′-UTR, a widely accepted hallmark of the m^6^A epitranscriptome. In this study, we selectively inhibited the helicase and ATPase activity of EIF4A3 and showed that these functions are critical to shaping the m^6^A landscape.

Xiang *et al*. reported a subtle anti-correlation between A-to-I editing and m^6^A in HEK cells by using MeRIP to capture all m^6^A-containing RNA transcripts and deep sequencing of the m^6^A-depeleted RNA in the supernatant (Jian-Feng Xiang et al., 2018). Consistent with our study, knockdown of METTL3 or METTL14 did not alter the number of inosine sites, however they observed enhanced and decreased A-to-I editing rates in about a third of the loci, with a slight excess of the former. Because EpiPlex enriches inosine sites, the A-to-G mutation rate under enrichment peaks does not correspond to the biological editing rate. While this makes EpiPlex more sensitive and allows for the detection of inosine in introns, a simple comparison between our measurements and Xiang’s A-to-G mutation-based editing rate cannot be made.

The EpiPlex proximity barcoding assay is a new approach to RNA modification detection that is inherently scalable to many modifications, pending the availability of specific RNA modification binders. Using densely modified *in vitro* transcribed RNA as target and antibody-loaded assay beads, the assay has been demonstrated for the simultaneous detection of five different RNA modifications (not shown). However, owing to the avidity effect, some antibodies require clusters of RNA modifications for specific binding, as is present in IVT RNA but not always in biological RNA. Due to this limitation, antibodies have limited merit for detecting rare RNA marks that occur at isolated sites. We are addressing this challenge by evolving new protein binders with superior specificity for biological RNA, which will be incorporated in future versions of the EpiPlex assay.

Our proximity barcoding approach offers a path to recording sequence-resolved RNA modification landscapes with high accuracy. A comparison with published m^6^A sequencing methods revealed that EpiPlex m^6^A data track closely with GLORI data (**SI FIG. 1**). In diagnostics, larger numbers of biomarkers are typically correlated with higher test accuracy. The multiplexing capability combined with relative quantification based on spike-in controls makes the EpiPlex assay ideal for biomarker discovery, validation and test development.

Today, the only other emerging tool for reading multiple RNA modifications are nanopores. Those have been proven particularly useful for studying abundant RNA species with high modification stoichiometries at conserved sites (tRNA, rRNA). This year, Oxford nanopore released a new pore and algorithm that together show promise for m^6^A and pseudouridine detection in mRNA but the signal-to-noise, need for an unmodified reference and 200 ng of mRNA per sample remain a challenge. The EpiPlex assay has been demonstrated to provide concordant results with GLORI with as little input as 20 ng of mRNA and 250 ng of total RNA, and the provided analysis pipeline is user-friendly and easy to deploy. Benchmarking studies with nanopore are under way.

## Methods

### EpiPlex RNA assay

Epitranscriptomic profiling was conducted using AlidaBio’s EpiPlex RNA library preparation kit (#EP100100) using 50 ng of purified, polyA-selected RNA as input. Poly-A selection was performed using the NEBNext® Poly(A) mRNA Magnetic Isolation Module (E7490L). In brief, purified RNA is mixed with spike-in controls, fragmented to ∼ 140bp, end-repaired and ligated to the P7 adapter. RNA fragments are captured by specific RNA modification binders. The captured RNA fragments are random primed by bead-anchored P5 adapters. The immobilized P5 adapters comprise the MBC (Modification Barcode), which is transferred to the cDNA during reverse transcription. PCR amplification concludes the library preparation and paired-end sequencing is used to read the UMI, MBC along with the RNA sequence. A solution control is processed in parallel, skipping the bead exposure. The EpiPlex kit contains all reagents needed to produce a library with Illumina adapters and unique dual sample indices. The libraries were QCed by Tapestation, quantitated by Picogreen, pooled and sequenced on a NextSeq 1000 in paired-end mode using 151,8,8,71 cycles. All libraries were sequenced at a coverage of at least 10 M reads. Since the writing of this manuscript, an updated reagent kit for m6A and inosine detection has become available (#EP100108).

### Data analysis

#### Peak Calling

Reads were processed by AMFIP (Alida Modification Finding Pipeline). Briefly, raw read quality was initially determined using FastQC (1). MBC and UMI barcodes were then extracted from read sequences, while low-quality bases were trimmed out. Trimmed reads with length <30bp were removed. Clean reads were then mapped to the human genome (GRCH38.p14) using STAR, as well as to our internal spike-in control, a modified Lambda Model System (LMS) references using Bowtie2. Genome mapped reads were split by MBC and deduplicated according to MBC and UMI barcodes using samtools (2) and a custom python script.

Peak calling was performed using a custom python script on a per-modification basis. Peaks were called in a signal-processing manner using the enrichment sample as signal and solution control as a measure of background. Peak regions and fold enrichment were determined using the Python package scipy (3), and were then scaled according to the LMS enrichment factor resulting in final peak fold-enrichment. The LMS enrichment factor was determined by scaling normalized LMS read counts on mod-specific LMS segments to an internal standard sequence containing a single mod with an internally measured standard curve (see SI Figs. 4-6). To identify high-confidence peaks, we employed a dynamic filtering technique based on the decay of fold-enrichment in false positive peaks as determined by reproducibility. Noise was modeled using an exponential decay curve with a linear component assuming a maximum false positive rate of 5 %. Samples that did not meet the 5% false positive rate criteria or failed the linearity assumption were defaulted to a fold-enrichment cutoff corresponding to the last cutoff where the linearity assumption held. Peaks which passed this cutoff, along with a minimum depth requirement of 5 reads in the enrichment sample, were considered to be “high-confidence”. After peak filtering, peaks were annotated using the GRCh38.p14 GTF and fasta files using a custom python script. Differential analysis was performed using bedtools and a custom python script. Gene clustering of peaks was performed using the Python package sklearn (4) AgglomerativeClustering. Aggregated z-scores were then clustered using Euclidean affinity and ‘ward’ linkage algorithms.

#### Variant Calling

To accurately identify A-to-G variants captured within EpiPlex inosine peak calls, we used the BCFtools (5) commands mpileup, norm, call, and filter with the mpileup parameters (--full-BAQ, --max-depth 20000), call parameters (-mA), and filter criteria (min(FMT/DP)>5)). Only sites which overlapped inosine peak calls and had a variant rate ≥10% were reported.

1) Andrews S. (2010). FastQC: a quality control tool for high throughput sequence data. Available online at: http://www.bioinformatics.babraham.ac.uk/projects/fastqc
2) Danecek P., Bonfield J. K., Liddle J., Marshall J., Ohan V., Pollard M. O., Whitwham A., Keane A., McCarthy S. A., Davies R. M., Li H.. Twelve years of SAMtools and BCFtools. GigaScience, Volume 10, Issue 2, February 2021, giab008, https://doi.org/10.1093/gigascience/giab008
3) Virtanen P., Gommers R., Oliphant T. E., Haberland M., Reddy T., Cournapeau D., Burovski E., Peterson P., Weckesser W., Bright J., van der Walt S. J., Brett M., Wilson J., Millman K. J., Mayorov N., Nelson A. R. J., Jones E., Kern R., Larson E., Carey C. J., Polat İ., Feng Y., Moore E. W., VanderPlas J., Laxalde D., Perktold J., Cimrman R., Henriksen I., Quintero E. A., Harris C. R., Archibald A. M., Ribeiro A. H., Pedregosa F., van Mulbregt P., and SciPy 1.0 Contributors. (2020) SciPy 1.0: Fundamental Algorithms for Scientific Computing in Python. Nature Methods, 17(3), 261-272, https://doi.org/10.1038/s41592-020-0772-5
4) Fabian Pedregosa, Gaël Varoquaux, Alexandre Gramfort, Vincent Michel, Bertrand Thirion, Olivier Grisel, Mathieu Blondel, Peter Prettenhofer, Ron Weiss, Vincent Dubourg, Jake Vanderplas, Alexandre Passos, David Cournapeau, Matthieu Brucher, Matthieu Perrot, Édouard Duchesnay; Scikit-learn: Machine Learning in Python, Pedregosa et al., JMLR 12, pp. 2825-2830, 2011.
5) Li H. A statistical framework for SNP calling, mutation discovery, association mapping and population genetical parameter estimation from sequencing data. Bioinformatics (2011) 27(21) 2987-93, https://doi.org/10.1093/bioinformatics/btr509

#### Mass spectrometry (HPLC-MS/MS) analysis of RNA

To be able to differentiate the de novo methylation from the existing methylation, HEK293T cells were incubated with media (DMEM + 10% FBS) containing heavy methionine (0.25 mg/mL CD_3_-Methionine (Cambridge Isotope, DLM-431-1) for 2 hours. Total RNA was isolated from the cells using TRIzol reagent. 3ug RNA was digested with 0.3 U P1 nuclease and dephosphorylated with 0.3 U rSAP at 37°C overnight. Then 100 μL samples were filtered with Millex-GV 0.22u filters.10 uL from each sample was injected into Agilent Triple Quad LC/MS instrument. The samples were run in mobile phase buffer A (water with 0.1% Formic Acid) and 2 to 20% gradient of buffer B (Methanol with 0.1% Formic Acid). MRM transitions were measured for adenosine (268.1 to 136.1), N6-methyladenosine (m^6^A) (282.1 to 150.1), and CD_3_-m^6^A (285 to 153). For LC/MS-MS data collection and analysis the following software was used: Agilent Mass Hunter LC/MS Data Acquisition Version B.08.00 and Quantitative Analysis Version B.07.01.

#### Generation of CRISPR KO cell lines

To generate knock-out cell lines for METTL3 and ADAR genes we used lentiCRISPR-V2 plasmid (lentiCRISPR v2 was a gift from Feng Zhang (Addgene plasmid # 52961; http://n2t.net/addgene:52961; RRID:Addgene_52961)).

The following primer sequences were used to clone guideRNAs into this vector:

hMETTL3 g1 intron-exon forward CACCGATGACTGGTGGAACGAACCT

hMETTL3 g1 intron-exon reverse AAACAGGTTCGTTCCACCAGTCATC

hMETTL3 g2 intron-exon forward CACCGTATGCCCCACAGGTATGAAC

hMETTL3 g2 intron-exon reverse AAACGTTCATACCTGTGGGGCATAC

To generate N terminal dTAG METTL3 expressing HEK293T cells, the cells were transduced with virus made using the full length METTL3 cloned into Blasticidin resistance gene containing pLEX_305-N-dTAG plasmid. (pLEX_305-N-dTAG was a gift from James Bradner & Behnam Nabet (Addgene plasmid # 91797; http://n2t.net/addgene:91797; RRID:Addgene_91797)). N term dTAG METTL3 expressing cells were selected with 10 ug/mL blasticidin for 3 days. Then the cells were transfected with guide RNAs targeting endogenous METTL3 and selected with 1 ug/mL puromycin for 3 days. Surviving cells were sorted as single cells into 96 well plates. Grown colonies were tested for knock-out of endogenous METTL3 and expression of dTAG-METTL3 via western blotting employing METTL3 antibody (Cell Signaling E3F2A #86132) and HA antibody (Santa Cruz F7 sc-7392).

To generate ADAR knock out HEK293T cells, the following primer sequences were used to clone guide RNAs into lentiCRISPR v2 plasmid:

hADAR g1 for CACCGGTCTCTTACCGAGGTTCATG

hADAR g1 rev AAACCATGAACCTCGGTAAGAGACC

hADAR g2 for CACCGGTCGTCGTCAGCTTGGGAAC

hADAR g2 rev AAACGTTCCCAAGCTGACGACGACC

ADAR knock out HEK293T cells were generated following the above methodology. ADAR knock-out for both p150 and p110 isoforms was confirmed by western blots employing ADAR antibody (Cell Signaling E6X9R XP #81284).

## Data availability

The raw FASTQ files have been deposited in the GEO database (NCBI), together with the m^6^A and inosine peak lists and bam files with the locations of A-to-G mutations. Summary statistics with the sequencing metric and peak counts are available as supporting info.

Link: https://www.ncbi.nlm.nih.gov/geo/query/acc.cgi?acc=GSE278242

## Code availability

The code that was used for data analysis is available upon request.

## Statistics and reproducibility

All experiments included two technical replicates of two biological replicates. Thus, error bars of peak numbers present the standard deviation of four data points. For the METTL3 inhibitor titration, the technical replicates were pooled to increase the read coverage and then downsampled to the same coverage at an MBC level. R^2^ values of differential fold-enrichment plots were calculated using standard Pearson correlation. For statistical testing we used a Wilcoxon rank-sum test where applicable.

### Acknowledgments

R.I.G. was supported by an Outstanding Investigator Award (R35CA232115) from the National Cancer Institute (NCI) of the NIH. Alida Biosciences was supported by Small Business Innovation Awards (R43 & R44 HG012170) from the National Human Genome Research Institute (NHGRI) of the NIH.

## Author contributions

E. S. and R. I. G. designed the experiments, and E.S. and J.K. executed them. H. Y., Y. H., J. S., and J. N. developed aspects of the EpiPlex assay. Q.L., A. A., and Z. M engineered & evolved protein binders. Z. J., A. C., Z. D., T. B. and. E. D. developed the analysis pipeline. A. P., B. W. P. and G.S. invented foundational concepts of the EpiPlex assay. G. S., E. S., R. I. G., Y. H. and Z. J. wrote the manuscript and prepared the figures. Z. M., A. P., G. S. and R. I. G. shared in the intellectual supervision of the work.

## Supporting Information

The Supporting Information is available free of charge at [URL].

- Supplementary figures.

