## Supporting Information for "Mapping multiple RNA modifications simultaneously by proximity barcode sequencing"

*Corresponding authors:

Richard I. Gregory

**SI FIG. 1**

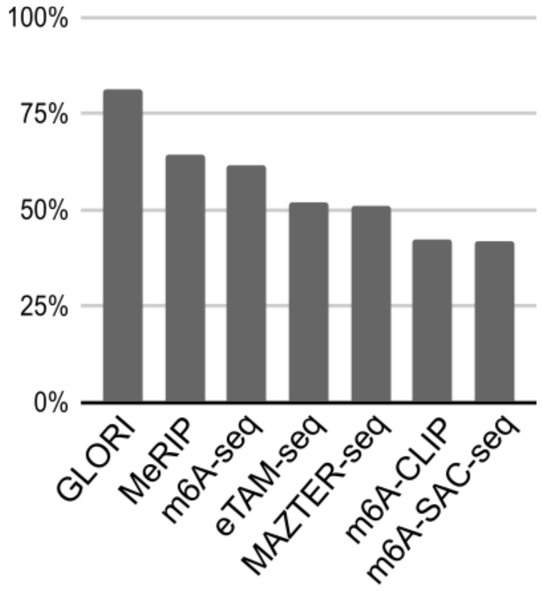


**SI FIG. 1. Comparison of EpiPlex $m^6A$ sites with public datasets.** Shown are the percentages of $m^6A$ peaks from EpiPlex overlapping with published datasets from GLORI[1] (HEK293T), meRIP[2] (HEK293T), $m^6A$-seq[3] (A549), eTAM-seq[4] (HeLa), MAZTER-seq[5] (HEK293T), $M^6A$-CLIP[6] (A549), and $m^6A$-SAC-seq[7] (HeLa). The EpiPlex $m^6A$ peaks used in this comparison are from DMSO treated HEK293T. The peaks are filtered by the quality score to limit the comparison to high confidence peaks.

Public data sets for comparison:

**SI FIG. 2A**

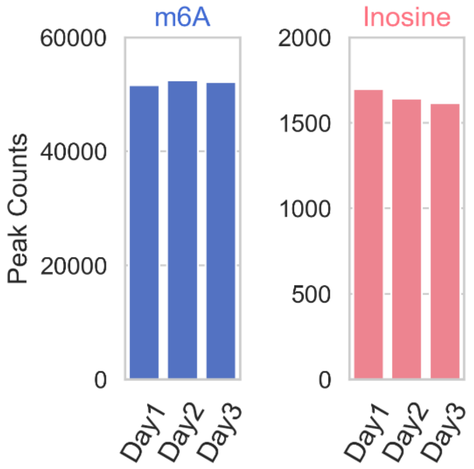


**SI FIG. 2A: Assay reproducibility across experiments.** Shown is the reproducibility of EpiPlex on the same sample across different days. The same DMSO treated and polyA enriched HEK293 RNA was used in three EpiPlex experiments on different days. The resulting peak counts for $m^6A$ and Inosine are highly consistent, with percent standard deviations at 0.80% and 2.55%, respectively.

**SI FIG. 2B**

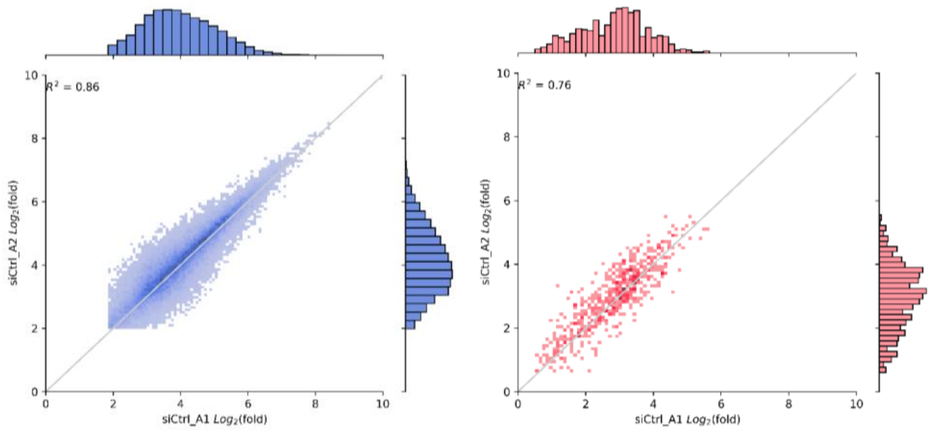


**SI FIG. 2B: Differential fold-enrichment across technical replicates.** Shown is the reproducibility of two technical replicates included in the same experiment. For $m^6A$, the $R^2$ is typically > 0.8 (blue) and the inosine $R^2$ is > 0.7.

SI FIG. 3

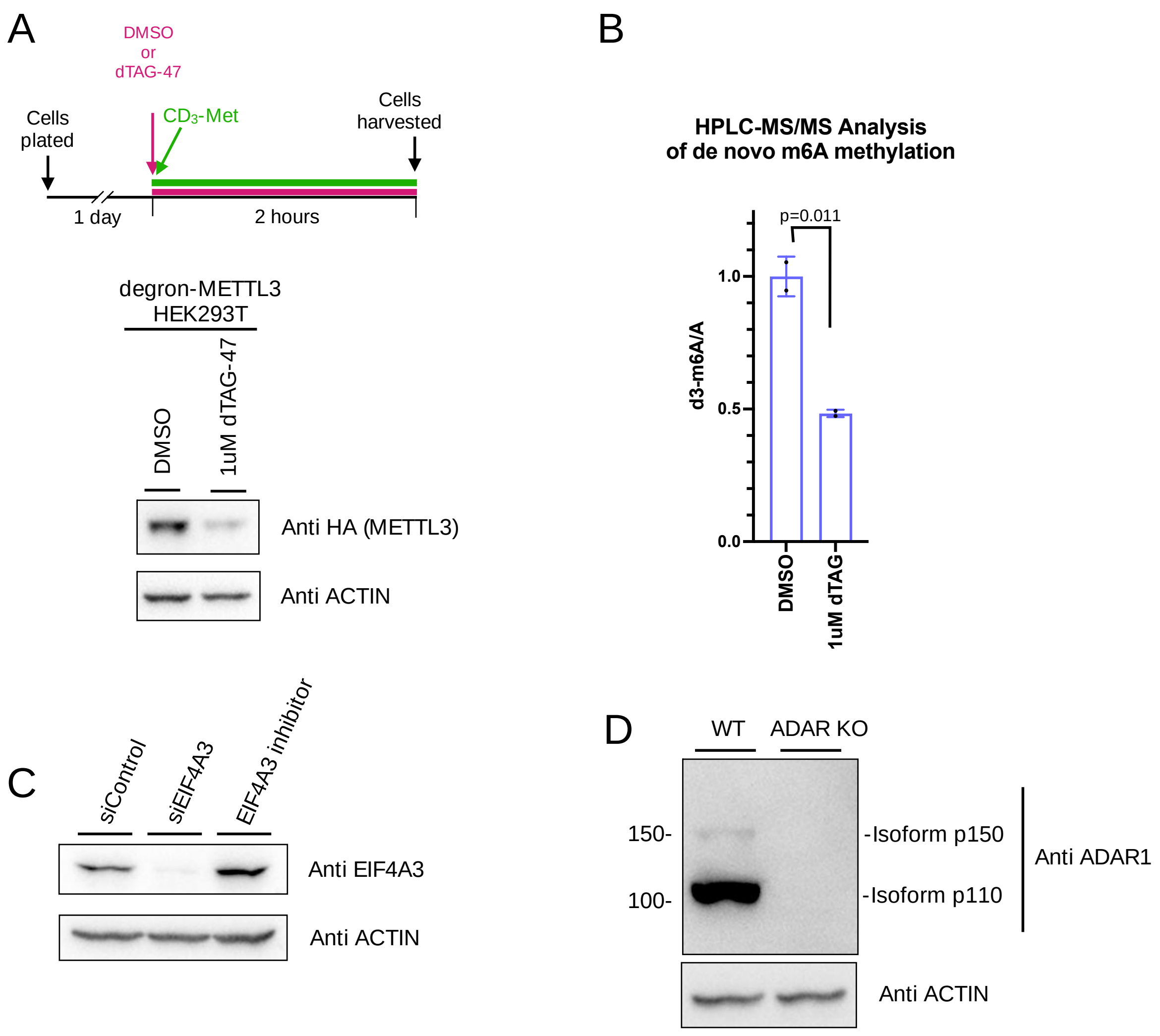


**SI FIG. 3: Degron mediated loss of METTL3 enzyme decreases *de novo* m$^6$A. (A)** Western blots indicating METTL3 levels upon degradation of METTL3 with dTAG treatment. **(B)** HPLC-MS/MS analysis of de novo m$^6$A (metabolically labelled $CD_3$-m$^6$A) that occurs during the 2 hour window in the presence of $CD_3$-Methionine and DMSO or 1 uM dTAG treatment. **(C)** Western blots indicating EIF4A3 levels upon siRNA or inhibitor treatment. **(D)** Western blots of control and ADAR1 KO HEK293T cells

**LMS Spike-In control**

The Lambda Model System (LMS) spike-in control is made from multiple *in vitro* transcribed fragments from the Lambda genome. Each fragment is around 1500 bases long, mimicking general mRNA sizes. The LMS is composed of several fragments without any modifications as a background, one set of fragments with $m^6A$ at high and low densities, and one set of fragments with Inosine at high and low densities. The modifications are incorporated into the fragments by adding modified nucleotides during *in vitro* transcription. The actual densities of the modifications are measured by HPLC.

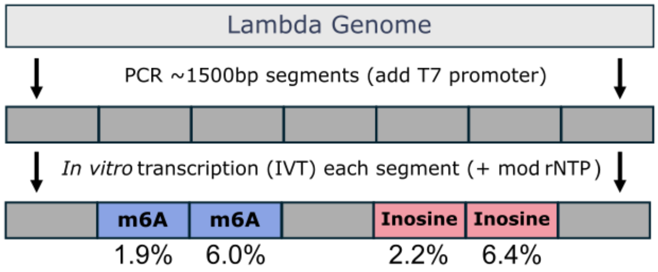


**SI FIG. 4: Lambda Model System (LMS) schematics.** The percentages indicate the amount of RNA mods determined for each fragment using HPLC.

**Enrichment Normalization**

For each modification barcode, ***mbc***, we first calculate the raw enrichment values of the sample ($E_{sample}(b_i)^{mbc}_{raw}$) at each base ***bi***, by dividing the depth-normalized coverage (***R$_{sample}$(bi)***, reads per million, rpm) at each base in the enrichment (***enrich***) over that in the solution control (***control***) (Eq.1). Note that the depth-normalized coverage for the solution control uses reads merging from all modification barcodes, where $R_{sample}(b_i)^{merged}_{control} = \sum_{mbc} R_{sample}(b_i)^{mbc}_{control}$.

$$E_{sample}(b_i)^{mbc}_{raw} = \frac{R_{sample}(b_i)^{mbc}_{enrich}}{R_{sample}(b_i)^{merged}_{control}} \qquad \textbf{Eq.1}$$

We then calculate the enrichment value for the LMS spike-in control similarly as the sample enrichment value in Eq.1. The difference is that we sum up all the reads aligned to each spike-in fragment to calculate fragment enrichment ($E_{lms}(f_i)^{mbc}_{raw}$, Eq.2), instead of base-by-base enrichment in Eq.1 to account for the random distribution of RNA mods across the fragments.

$$E_{lms}(f_i)^{mbc}_{raw} = \frac{\sum_i R_{lms}(b_i)^{mbc}_{enrich}}{\sum_i R_{lms}(b_i)^{merged}_{control}}$$ **Eq.2**

The sample raw enrichment values for each ***mbc*** are then normalized by dividing by the raw enrichment of the standard fragment $\boldsymbol{f_{mbc}}$ in the spike-in control and multiplying by a scaling factor, $\boldsymbol{N^{mbc}}$. $\boldsymbol{N^{mbc}}$ has been optimized to account for the different enrichment strengths of our multiplexing binders.

$$E_{sample}(b_i)^{mbc}_{norm} = E_{sample}(b_i)^{mbc}_{raw} \times \frac{N^{mbc}}{E_{lms}(f_{mbc})^{mbc}_{raw}}$$ **Eq.3**

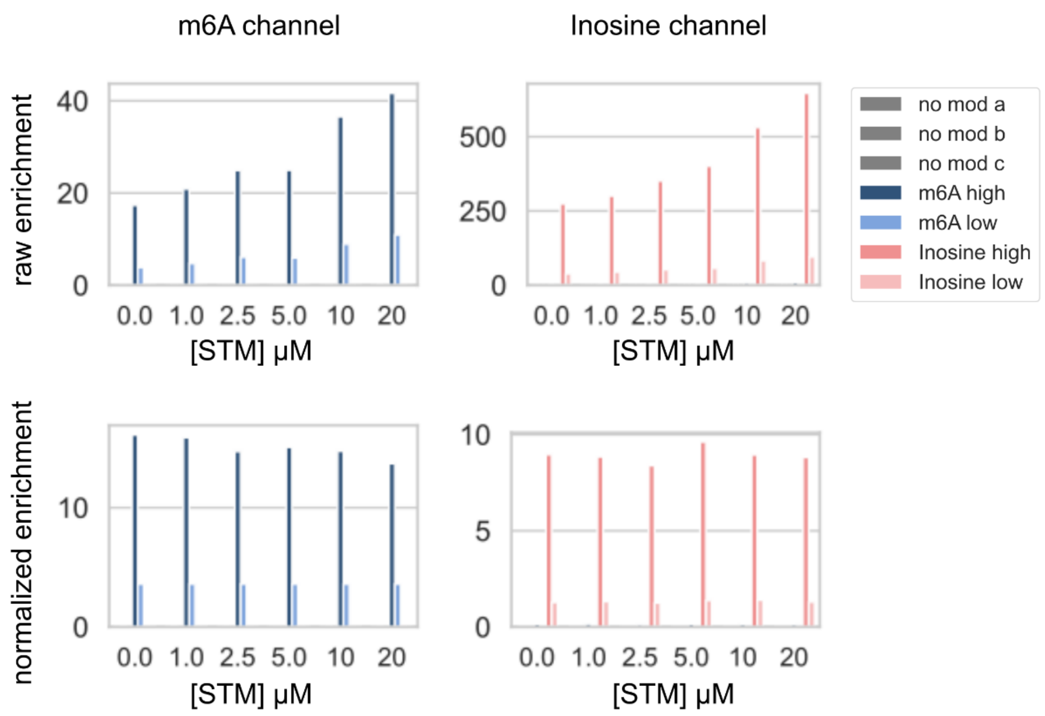


**SI FIG. 5: Signal normalization.** The plots demonstrate how our LMS spike-in control calibrates the signal from systematic drifts due to biological effects in the sample. This is important to enable cross-sample comparisons. In the STM titration experiment, the $m^6A$ sites in the sample reduce as the concentration of STM increase. Consequently, the remaining $m^6A$ sites in both the sample and the LMS receive higher raw enrichment values as less competition for the binding sites and sequencing. This phenomenon is reflected in the LMS spike-in control for both $m^6A$ and Inosine channels (top row, raw enrichment). Since the modification fragments in LMS spike-in control have constant modification sites in all STM concentrations, they serve as an anchor point to the real enrichment value. After the operation by Eq.3, the normalized enrichment values return to a stable value across all STM concentrations (bottom row, normalized enrichment).

**SI FIG. 6**

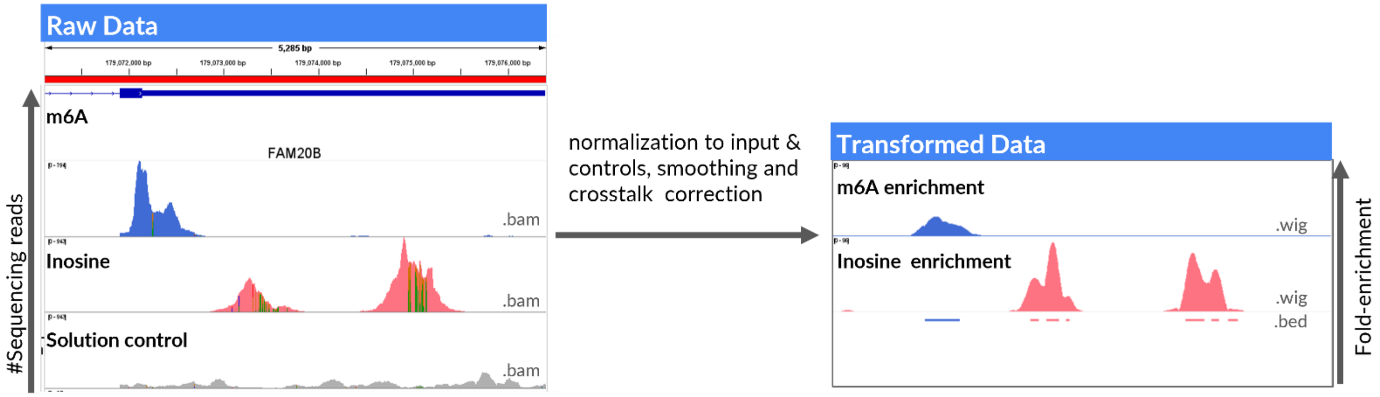


**SI FIG. 6: Data transformation prior to peak calling.** The left panel shows the bam files, representing the raw reads of the $m^6A$ enrichment reaction (blue), the inosine enrichment reaction (red – the vertical lines indicate A-to-G mutations) and the solution control (gray). The right panel shows the result of the data transformation as wig files. The horizontal bars under the peaks are the bed files, indicating the peak summits.
